# Cell cycle checkpoints cooperate to suppress DNA and RNA associated molecular pattern recognition and anti-tumor immune responses

**DOI:** 10.1101/2020.06.24.168971

**Authors:** Jie Chen, Shane M Harding, Ramakrishnan Natesan, Lei Tian, Joseph L Benci, Weihua Li, Andy J Minn, Irfan A Asangani, Roger A Greenberg

## Abstract

The DNA dependent pattern recognition receptor, cGAS mediates communication between genotoxic stress and the immune system. Mitotic chromosome missegregation is an established stimulator of cGAS activity, however, it is unclear if progression through mitosis is required for cancer cell intrinsic activation of immune mediated anti-tumor responses. Moreover, it is unknown if disruption of cell cycle checkpoints can restore responses in cancer cells that are recalcitrant to DNA damage induced inflammation. Here we demonstrate that prolonged cell cycle arrest at the G2-mitosis boundary from either CDK1 inhibition or excessive DNA damage prevents inflammatory stimulated gene expression and immune mediated destruction of distal tumors. Remarkably, DNA damage induced inflammatory signaling is restored in a cGAS-and RIG-I-dependent manner upon concomitant disruption of p53 and the G2 checkpoint. These findings link aberrant cell progression and p53 loss to an expanded spectrum of damage associated molecular pattern recognition and have implications for the design of rational approaches to augment antitumor immune responses.

## Introduction

Emerging evidence indicates that the efficacy of radio- and chemo- therapies requires DNA damage induced activation of cytotoxic immune responses (Formenti et al., 2018; Lee et al., 2009; Liang et al., 2013; Postow et al., 2012). The underlying mechanism for how radiotherapy activates anti-tumor immune responses remains obscure but is thought to involve radiation stimulated expression of type I interferon and other cytokines in cancer cells and surrounding stroma (Burnette et al., 2011; Deng et al., 2014; Woo et al., 2014). Studies from our lab and others demonstrate mitotic progression following genotoxic stress is required to activate type I interferon signaling that is associated with the pattern recognition receptor (PRR) cGAS localizing to cytosolic DNA within micronuclei (Bakhoum et al., 2018; Harding et al., 2017; Mackenzie et al., 2017; Santaguida et al., 2017; Yang et al., 2017) Whether cell cycle progression impacts the efficacy of combined DNA damaging and immune therapies remains unknown. Similarly, it is unclear whether activation of such responses is feasible in cells that demonstrate persistent cell cycle arrest or loss of the cGAS-STING pathway.

Irradiation (IR)-induced DNA damage responses invoke double-strand break (DSB) repair and cell cycle checkpoints that delay entry into S phase or mitosis. Such events are thought to allow adequate time for DSB repair. The IR-induced G2/M cell cycle checkpoint requires the Ataxia Telangiectasia and Rad3 related (ATR) – Checkpoint kinase 1(CHK1) pathway with additional contributions from the Ataxia Telangiectasia Mutated (ATM) kinase (Abraham, 2001; Liu et al., 2000; Xu et al., 2002). The G1/S cell cycle checkpoint is dependent on ATM mediated p53 induction and transcriptional activation of its target genes (Barlow et al., 1997; Canman et al., 1998; Kastan et al., 1992). Tumor cells with unstable genomes more frequently missegregate chromosomes during mitosis, leading to the formation of micronuclei. Nuclear envelope integrity is compromised in approximately 50% of micronuclei, allowing cGAS and other cytoplasmic proteins to recognize their dsDNA contents (Hatch, 2018; Hatch et al., 2013; Liu et al., 2018). This localization correlates with cGAS-production of its enzymatic product cyclic GMP-AMP (cGAMP) and subsequent activation of its signal transducer STING in cells that have experienced genotoxic stress (Coquel et al., 2018; Dou et al., 2017; Gluck et al., 2017; Harding et al., 2017; Mackenzie et al., 2017). The detection of foreign cytosolic RNA is mediated largely by RIG-I-like receptors (RLRs) including RIG-I, MDA5 and LGP2 (Ablasser and Hur, 2020). RNA polymerase III dependent transcription on cytosolic DNA has also been reported to stimulate RIG-I dependent inflammatory cytokine production (Ablasser et al., 2009; Chiu et al., 2009).

Recent findings illuminate several distinct possibilities to limit inflammatory responses to genotoxic agents. We reported that deficiency in canonical non-homologous end joining (c-NHEJ) abrogates micronuclei formation and renders cells unable to activate cGAS-STING dependent inflammatory signaling in response to IR-induced DNA damage (Harding et al., 2017). Tumor cells can also escape immune surveillance by silencing the cytosolic DNA-sensing pathway (Kwon and Bakhoum, 2020), preventing signaling responses to inflammatory cytokines, or suppression of antigenic peptide presentation (Benci et al., 2019; Ishizuka et al., 2019; Patel and Minn, 2018). cGAS or STING expression is reported to be reduced in many cancer cell lines, including melanoma, and in tumor cells that rely on alternative telomere maintenance, or express oncogenic DNA tumor viruses (Chen et al., 2017; Lau et al., 2015; Xia et al., 2016). These clinically relevant obstacles necessitate alternative approaches that can abrogate persistent cell cycle checkpoint activation and promote inflammatory signaling irrespective of canonical DNA sensing by cGAS-STING. Here we delineate the importance of DNA damage induced cell cycle checkpoints in relation to anti-tumor immune responses and describe cooperation between ATR and p53 dependent cell cycle checkpoints in limiting activation of DNA and RNA sensing pattern recognition receptors.

## Results

### CDK1 or c-NHEJ inhibition suppresses DNA damage-induced inflammatory signaling and anti-tumor immune responses

We performed a kinetic analysis of gene expression to decipher the complexity of inflammatory signaling pathways that are activated in response to ionizing radiation (IR) induced DNA damage. MCF10A cells were irradiated with 10 Gy and collected for RNA-seq analysis at three and five days, respectively (Figure 1A). Gene Set Enrichment Analysis (GSEA) (Subramanian et al., 2005) revealed that interferon alpha, interferon gamma and IL6-JAK-STAT3 signaling were the top activated inflammatory pathways following IR treatment (Figure 1B). Interestingly, interferon alpha and interferon gamma signaling further increased at five days compared to three days post IR, however, IL6-JAK-STAT3 signaling plateaued from day three onwards, suggesting these pathways might be regulated differently. Next, we asked if either of each inflammatory signature would be affected if cell cycle arrest was achieved to prevent entry to mitosis. Indeed, the CDK1 specific inhibitor RO-3306 (CDK1i) prevented the activation of all three signaling pathways (Figure 1C). Interestingly, c-NHEJ deficiency by loss of XRCC4 (Rooney et al., 2004) also significantly compromised inflammatory signaling (Figure 1D). XRCC4 is not required for Ku70/Ku80 DSB end recognition or activation of DNA-PKcs, but rather for the final ligation step of c-NHEJ (Graham et al., 2016). We therefore postulated that reductions in interferon stimulated gene expression are secondary to cell cycle arrest prior to mitosis due to excessive unrepaired DNA damage and not to a direct role in inflammatory signaling. Indeed, we observed persistent DNA damage indicated by γH2AX in XRCC4 knockout (KO) cells even 3 days post 10 Gy IR (Figure S1A). In contrast to wild type (WT) cells that initially arrested at G2 and began to progress through mitosis 16 h after IR, XRCC4 deficient cells remained persistently arrested at the G2-mitosis boundary (Figure S1B). In contrast, a low dose of 2 Gy irradiation that induced less DNA damage restored micronuclei formation and inflammatory signaling in MCF10A XRCC4 KO cells (Figures S1C and S1D). This observation is consistent with a model whereby c-NHEJ deficiency impairs cell cycle progression as a consequence of excessive DNA damage after 10 Gy rather than through the putative role of c-NHEJ factors (i.e. DNA-PK) in pattern recognition (Burleigh et al., 2020; Ferguson et al., 2012; Zhang et al., 2011). We further analyzed inflammatory gene expression in genetically defined MCF10A cells. IR treatment stimulated 81 differentially expressed genes, the majority of which were up-regulated (Figure 1E). Loss of pattern recognition receptor cGAS or STING compromised this increase, confirming their roles in mediating IR-induced inflammatory signaling. Notably, CDK1i treated cells suppressed these inflammatory signatures to a greater extent, suggesting that progression through mitosis is the more critical determinant in the activation of IR-induced inflammatory signatures.

**Fig 1.**
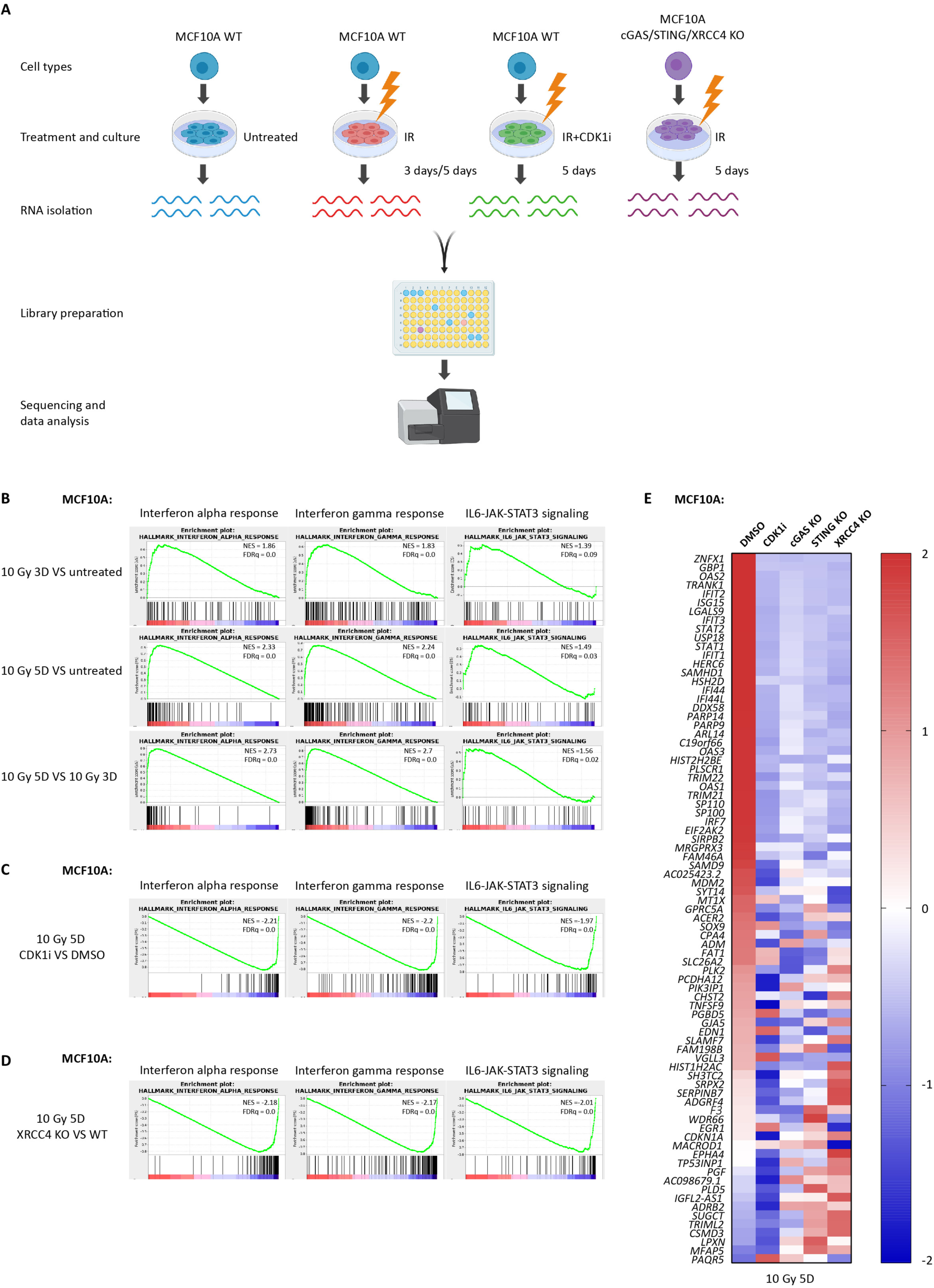
G2/M cell cycle arrest or c-NHEJ deficiency suppresses IR-induced inflammatory signaling. (**A**) Scheme showing the library preparation workflow for RNA-seq. Created with BioRender. (**B, C and D**) Gene set enrichment analysis (GSEA) of RNA-seq data in MCF10A cells to identify enriched biological pathways at 3 days after 10 Gy vs untreated, 5 days after 10 Gy vs untreated, and 5 days vs 3 days after 10 Gy (**B**); 5 days after 10 Gy with CDK1i vs 5 days after 10 Gy with DMSO (**C**); and 5 days after 10 Gy in XRCC4 KO vs 5 days after 10 Gy in WT (D), respectively. Significant GSEA Enrichment Score curves were noted for interferon alpha response, interferon gamma response, and IL6-JAK-STAT3 signaling. The green curve in the displayed GSEA thumbnails represents the enrichment score curve. Genes on the far left (red) correlated with treatment condition, and genes on the far right (blue) correlated with control condition. The vertical black lines indicate the position of each gene in the studied gene set. The normalized enrichment score (NES) and false discovery rate (FDRq) are shown for each pathway. (E) Heatmap showing the z-score of FPKM expressions in control, CDK1i, cGAS KO, STING KO, and XRCC4 KO MCF10A cells for genes differentially expressed at 5 days after 10 Gy treatment (fold change > 2 and p value < 1e-10).

Given the necessity of mitotic progression in DNA damage activated inflammatory signaling, we speculated that progression through mitosis might also play a critical role in anti-tumor immunity. To this end, we employed the B16F10 murine melanoma model to determine whether cell cycle arrest or c-NHEJ deficiency affected anti-tumor immunity (Figure 2A). Untreated B16 WT cells, were subcutaneously injected in one flank of C57BL/6J mice two days prior to injection of *ex vivo* treated B16 cells in the opposite flank. This model provides a method to track anti-tumor immunity without the confounding effect of systemic administration of drugs to animals (e.g. CDK1i) that may have unanticipated consequences including on the immune system. Importantly, we have previously shown that *ex vivo* irradiation and subsequent implantation of tumor cells produced equivalent responses to *in vivo* radiation as is classically done in abscopal models (Harding et al., 2017). Moreover, that anti-CTLA4 alone had limited impact on the abscopal tumors but addition of radiotherapy markedly increased this response (Harding et al., 2017; Twyman-Saint Victor et al., 2015). The different treatments of the primary tumor included 10 Gy IR, CDK1i or CRISPR-Cas9 mediated deletion of the c-NHEJ gene *Ku70* (Ku70 KO). Immune checkpoint blockade was achieved using anti-CTLA-4 (clone 9H10) antibody administered by intraperitoneal injection every three days for a total of three doses as previously described (Harding et al., 2017). Growth rates of untreated tumors were then monitored following the final anti-CTLA-4 antibody injection. Consistent with suppression of inflammatory signatures in RNA-seq data in Figure 1E, cell cycle arrest by CDK1i abolished the IR-induced anti-tumor effect (Figures 2B and 2D). These data agree with the requirement for mitotic progression in irradiated tumor cells to activate systemic immune responses. We considered an alternative possibility that an extended G2 arrest during CDK1i treatment may enhance DNA repair in these cells, thus accounting for the reduction in immune activation following IR. To address this issue, we examined how loss of the critical c-NHEJ repair factor Ku70 would affect systemic immune responses. Interestingly, Ku70 deleted B16F10 cells also led to a failure of anti-tumor immune responses to limit the growth of untreated tumors (Figures 2C and 2E) despite the importance of c-NHEJ in repairing IR-induced DNA damage. Taken together, these findings indicate that progression through mitosis is a key gatekeeper for IR-induced inflammatory signaling and anti-tumor immunity.

**Fig 2.**
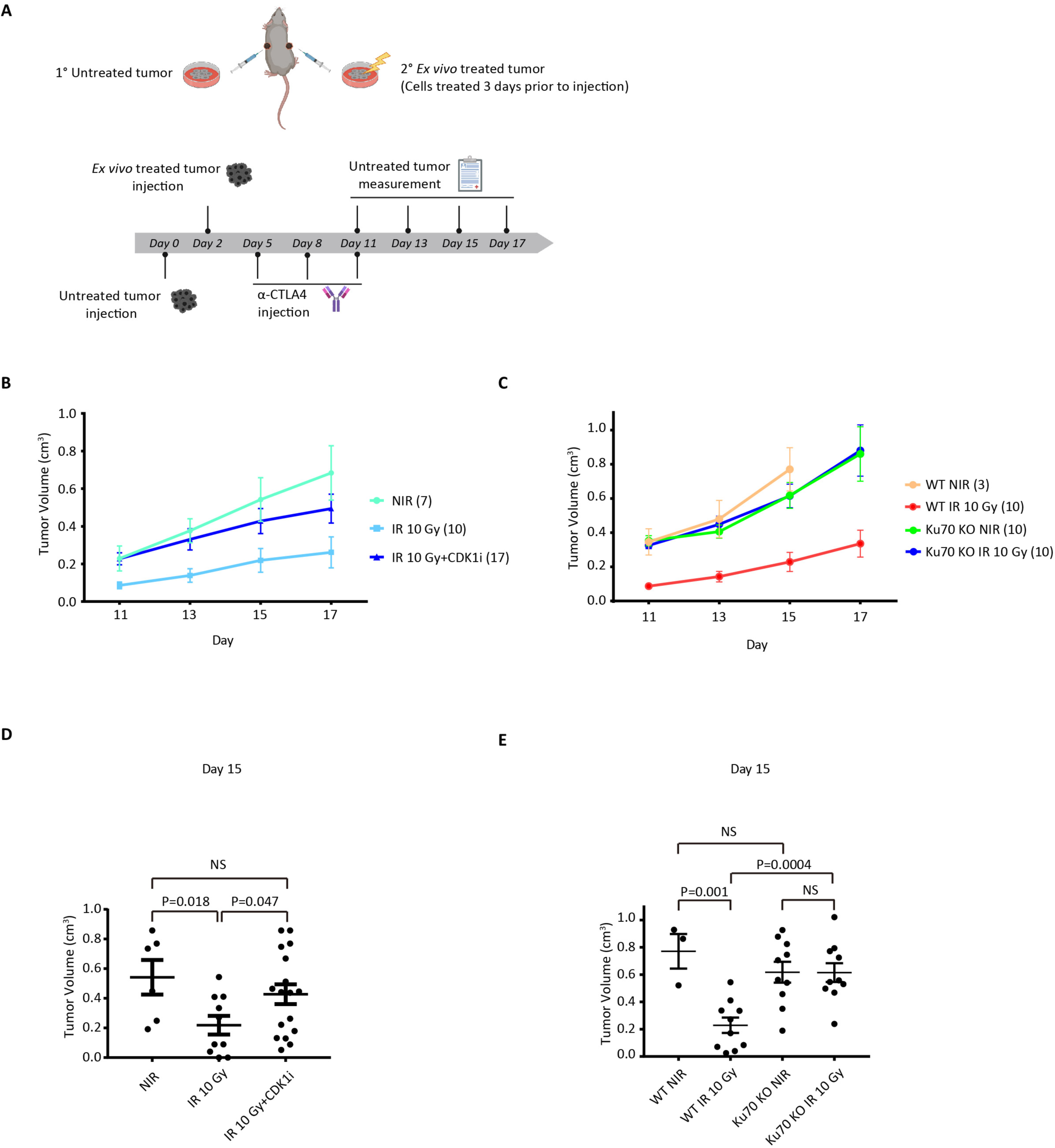
Progression of the irradiated tumor cells through mitosis is required for systemic anti-tumor immune responses. (**A**) Scheme showing B16 melanoma working model. Created with BioRender. (**B and C**) Growth of wild type B16 cells (untreated tumors) following injection of B16 wild type cells with no treatment, 10 Gy irradiation, or 10 Gy irradiation with CDK1i treatment (B), and wild type B16 cells or Ku70 KO B16 cells with no treatment or 10 Gy irradiation (**C**) 3 days before implantation, respectively. All mice were administrated with anti-CTLA-4 antibody (9H10) as described in **A**. (**D and E**) Statistic of tumor volumes at day 15 as measured in **B** and **C**. Statistical significance is compared using two tailed t-test. Error bars are S.E.M of biological replicates.

### Cell cycle checkpoint abrogation accelerates micronuclei formation and inflammatory signaling

A corollary to the prior experiments is that abrogation of the G2/M checkpoint would enhance the kinetics and amplitude of micronuclei formation and inflammatory signaling. To test this possibility, we irradiated cells with 20 Gy, a dose expected to induce a strong G2/M cell cycle checkpoint in combination with inhibitors of ATR or CHK1 kinase activity. Contrary to the limited micronuclei formation in DMSO treated cells, approximately 50% of ATR inhibitor (ATRi) or CHK1 inhibitor (CHK1i) treated cells formed micronuclei just 24 hours post 20 Gy IR. A considerable portion of cells formed micronuclei in the DMSO treated group 72 hours after IR, however, their frequency was still increased after ATRi or CHK1i treatment (Figure 3A). Cell cycle analysis by flow cytometry confirmed abrogation of the G2/M checkpoint by ATRi or CHK1i (Figure 3B). In response to 20 Gy irradiation, most cells arrested in G2. ATRi or CHK1i did not alter the cell cycle distribution in non-irradiated cells, but greatly increased G1 phase populations in irradiated cells, consistent with cells traversing mitosis. To further determine if cells progressed through mitosis following IR, we performed dye dilution experiments (Figure S1E) that enable quantification of cell division numbers over a given time interval. Reductions in fluorescence indicate cell division due to dye dilution. Irradiated MCF10A cells underwent 1-2 divisions over 3 days confirming mitotic progression after IR. This change in dye content was suppressed by CDK1i administration. ATRi/CHK1i treated cells exhibited evident STAT1 Y701 phosphorylation after 24 hours that was strongly increased at three days (Figure 3C). Prevention of mitotic entry using either CDK1i or a PLK inhibitor (PLKi) abolished the enhanced inflammatory signaling (Figure 3D).

**Fig 3.**
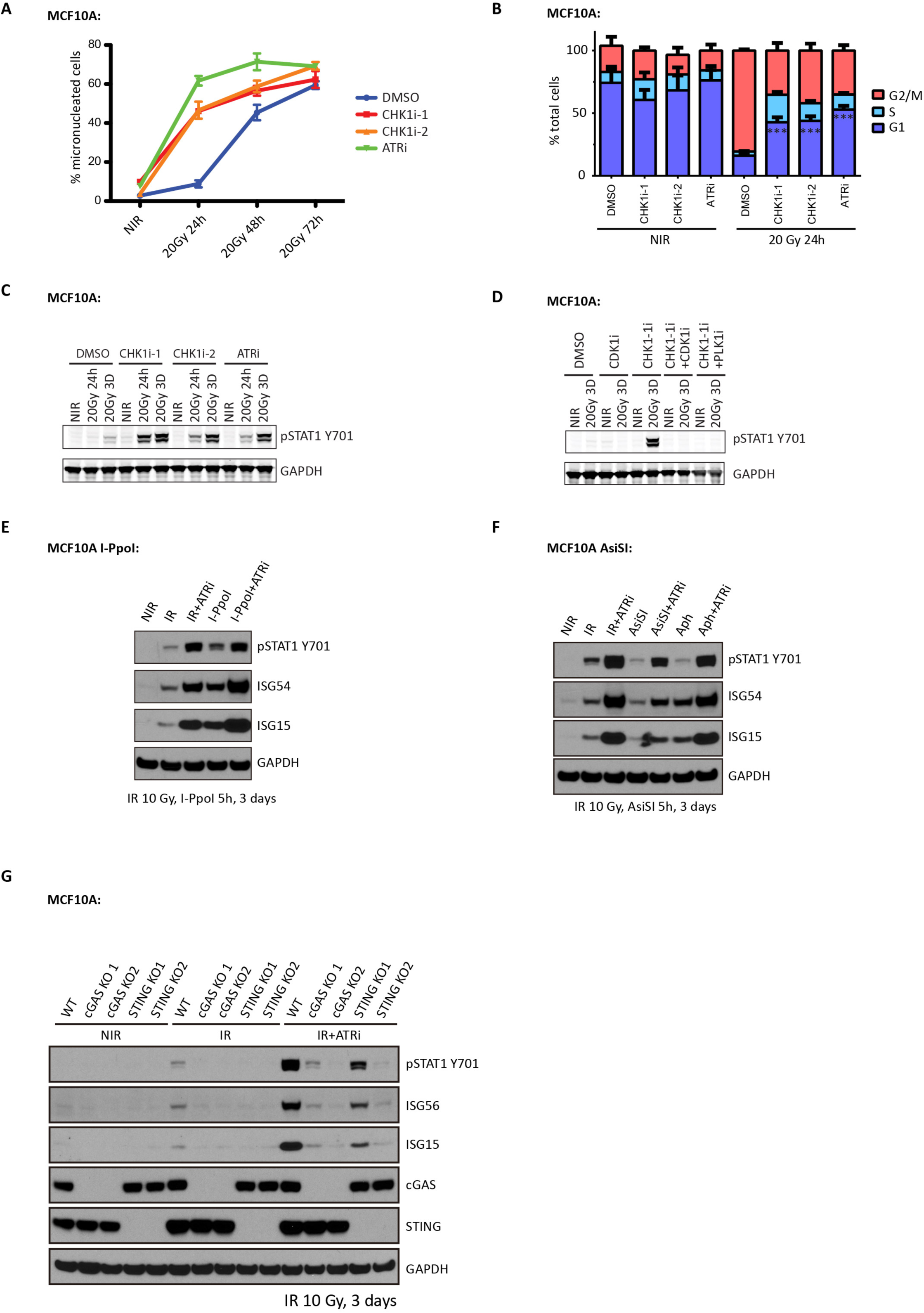
Disruption of the IR-induced G2/M checkpoint enhances inflammatory signaling activation in MCF10A cells. (**A**) MCF10A cells were irradiated or left untreated with or without the indicated treatments followed by fixation at the indicated times. Cells with micronuclei were quantified. (**B**) MCF10A cells irradiated or left untreated with or without indicated inhibitors were fixed and subjected to cell cycle analysis by flow cytometry. (**C and D**) MCF10A cells irradiated with indicated dose or left untreated in the presence of indicated inhibitors were collected at specified time point for western blot. (**E**) MCF10A I-PpoI cells were left untreated (NIR), irradiated with 10 Gy (IR), or treated with 4-OHT and shield-1 for 5 hours (I-PpoI), and then maintained in medium with or without ATR inhibitor for 3 days before collection for western blot analysis. (**F**) MCF10A AsiSI cells were left untreated (NIR), irradiated with 10 Gy (IR), induced with 4-OHT and shield-1 for 5 hours (AsiSI), or cultured in the presence of aphidicolin (Aph), and then maintained in medium with or without ATR inhibitor for 3 days before collection for western blot analysis. For Aph treated cells, 2.5 μM aphidicolin were in the medium all the time until cell collection 3 days later. (**G**) MCF10A cells left untreated or irradiated with 10 Gy were maintained for 3 days in the presence or absence of ATR inhibitor before collection for western blot analysis. Error bars represent S.E.M.

Consistently, ATR inhibitor treated cells following 10 Gy irradiation also showed increased inflammatory signaling (Figure 3 E-G), demonstrating disruption of the G2 checkpoint produces similar results across a range of damage induction. Moreover, the increased inflammatory signaling was not limited to IR-induced DNA damage. ATRi treatment also increased inflammatory signaling in response to DSBs induced by two different nucleases I-PpoI and AsiSI (Figures 3E and 3F) and in response to replication stress using aphidicolin (Figure 3F). Increased inflammatory signaling after ATRi was also largely dependent on cGAS-STING pathway (Figure 3G), further highlighting the importance of mitotic progression for activation of this DNA sensing axis.

### Disruption of G2 and G1 checkpoints cooperate to restore IR-induced immune responses in c-NHEJ deficient cells

The observation that G2 checkpoint disruption was able to promote micronuclei formation and inflammatory signaling encouraged us to test whether ATRi treatment can activate interferon stimulated gene expression in irradiated cells that lack c-NHEJ repair. Unexpectedly, inhibition of ATR failed to restore micronuclei formation after 10 Gy (Figures 4A and 4B), and only slightly increased inflammatory signaling in XRCC4 KO cells (Figures 4D and 4E). In contrast, ATRi worked in XRCC4 KO cells upon a low dose of irradiation at 2 Gy (Figures S1C and S1D). This minimal effect can be partially explained by mitotic progression in cells with excessive DNA damage leading to mitotic catastrophe (Vakifahmetoglu et al., 2008). Indeed, live cell imaging showed that 85% of XRCC4 KO cells died during mitosis upon releasing cells at the G2/M boundary following 10 Gy compared to only 30% of WT cells (Figures S2A and S2B).

**Fig 4.**
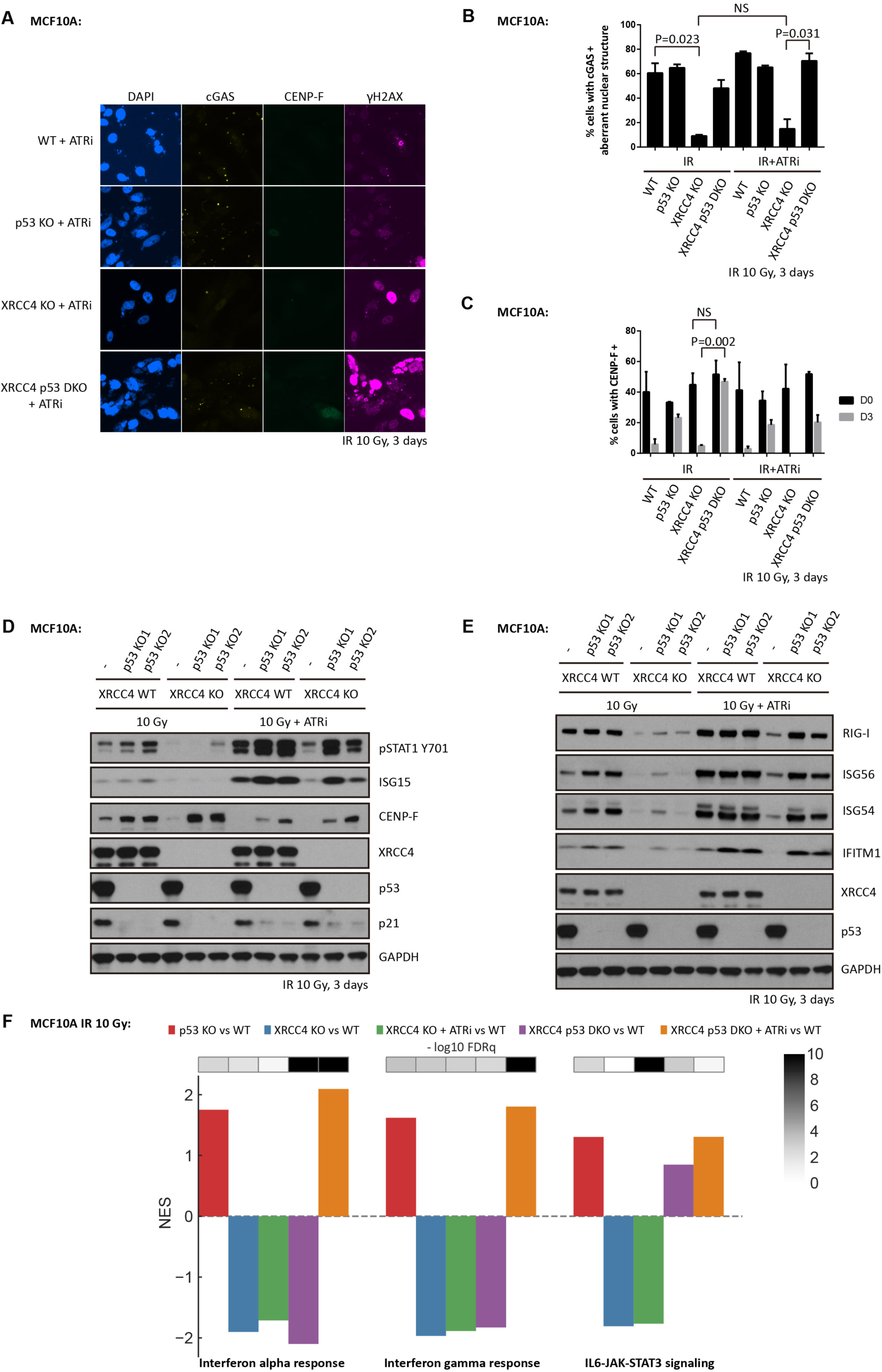
Disruption of p53 and ATR restores IR-induced inflammatory signaling in c-NHEJ deficient MCF10A cells. (A, B and C) WT cells, p53 KO cells, XRCC4 KO cells or XRCC4 P53 DKO cells were irradiated with 10 Gy (IR), and then maintained in medium with or without ATR inhibitor for 3 days before fixation for immunofluorescence staining. (D and E) WT cells, p53 KO cells, XRCC4 KO cells or XRCC4 P53 DKO cells were irradiated with 10 Gy and then cultured for 3 days in the presence or absence of ATR inhibitor before collection for western blot analysis. (F) Bar plot showing the normalized enrichment score (NES) and false discovery rate (FDRq) for the identified enriched biological pathways in MCF10A cells 3 days post 10 Gy irradiation. Error bars represent S.E.M.

To understand if mitotic progression after DSB induction is sufficient to activate inflammatory signaling, we released cells from G2 arrest following irradiation and examined inflammatory signaling by immunoblot. Surprisingly, direct release of irradiated cells from G2 resulted in minimal micronuclei formation and inflammatory signaling as assessed by ISG15 and STAT1 Y701 phosphorylation on immunoblot at three days post IR (Figures S2C and S2D). The cells that exhibited micronuclei in this population were EdU positive, suggesting cell cycle progression through S phase and a second round of mitosis had occurred (Figures S2C, S2E and S2F). This finding prompted us to ask whether the entry into S-phase with damage in MCF10A XRCC4 KO cells is also a critical barrier to inflammatory activation in addition to the G2/M checkpoint. Given the important role of ATM and p53 in the IR-induced G1/S checkpoint (Barlow et al., 1997; Canman et al., 1998; Kastan et al., 1992), we generated MCF10A p53 KO cell lines using CRISPR-Cas9 to test the importance of the G1/S checkpoint disruption in this inflammatory response to DNA damage. p53 loss led to increased S phase entry indicated by EdU incorporation (Figure S2E) and mildly higher percentages of cells with micronuclei (Figure 4A and 4B and Figure S2F). Despite equal cell death rates during mitosis, p53 loss increased cell viability in the subsequent interphase at 24 hours post mitosis (Figure S2G) consistent with its known role in responding to DNA damage (Nikoletopoulou et al., 2013). Introduction of S phase entry inhibitors in cells released from G2 resulted in minimal cell populations with micronuclei at three days after IR in WT cells (Figures S2E and S2F). However, there were much higher percentages of cells with micronuclei in p53 KO cells, which further suggested a role of p53 in cell death during the subsequent interphase following mitotic progression with DNA damage (Figures S2E and S2F).

The preceding findings suggested that several cell cycle checkpoints affect pattern recognition in response to genotoxic stress. We therefore generated MCF10A cells with deletion of both XRCC4 and p53 and subjected them to 10 Gy irradiation in the presence of ATRi to understand if combined loss of the G1/S and G2/M checkpoints could restore inflammatory signals. p53 loss increased the cell populations accumulated at S-and G2-phases as indicated by CENP-F positivity in cells at three days post IR (Figures 4C and 4D). Remarkably, p53 loss combined with ATRi restored micronuclei formation (Figures 4A and 4B) and inflammatory signaling (Figures 4D, 4E and 4F). Like p53 loss, ATM inhibition cooperated with ATRi to restore inflammatory signaling in XRCC4 deficient MCF10A cells (Figures S3A and S3B). Restored inflammatory signaling was also consistently found upon disruption of ATR and p53 checkpoints in Ku70 deficient cells (Figures S3C and S3D).

### ATR inhibition cooperates with p53 mutation to activate cGAS-independent inflammatory signaling

Loss of cGAS or STING did not completely eliminate inflammatory signaling upon ATRi treatment (Figure 3G), leading us to speculate the presence of additional damage associated molecular patterns. To address this possibility, we used hTERT immortalized retinal pigmented epithelial (RPE-1) cells, a cell line that shows undetectable cGAS expression by either immunofluorescence or western blot (Figures S4A and S4B). Similar levels of STING expression were present in RPE-1 cells when compared to MCF10A cells (Figure S4B). In response to irradiation and ATRi, RPE-1 WT cells failed to discernably activate inflammatory signaling (Figures 5A and 5B), despite approximately 50% of cells containing micronuclei (Figures 5C and 5D). Strongly increased inflammatory signaling occurred in RPE-1 WT cells upon ectopic cGAS expression but not following introduction of a cGAS inactive mutant K411A (Figures 5A and 5B), revealing that WT cGAS promoted signaling responses to missegregated genomic DNA in RPE-1 cells as it has been shown in MCF10A and many other independent cell lines. We also observed the RNA sensing pattern recognition receptor RIG-I was markedly increased in RPE WT cells with ectopic cGAS after IR, consistent with reports that RIG-I expression is inflammatory cytokine inducible (Liu and Gu, 2011; Yuzawa et al., 2008). Surprisingly, inflammatory signaling was strongly present in ATRi treated RPE-1 p53 KO cells in response to 10 Gy IR irrespective of cGAS (Figures 5B). Endogenous cGAS expression was not detected in either p53 WT or mutant RPE-1 cells (Figures 5A and 5B) arguing against its involvement in damage induced inflammatory signaling. Notably, inflammatory stimulated gene expression remained dependent on mitotic progression since CDK1i abolished this effect (Figure 5B). We performed RNA-seq analysis to independently confirm these results and more extensively compare gene expression in each of the different settings. Similar pathway activation occurred in both RPE-1 WT cells with ectopic cGAS expression and in RPE-1 p53 KO cells that lack cGAS expression, with the interferon gamma, interferon alpha and IL6-JAK-STAT3 signaling responses being most significantly elevated in each group (Figure 5E).

**Fig 5.**
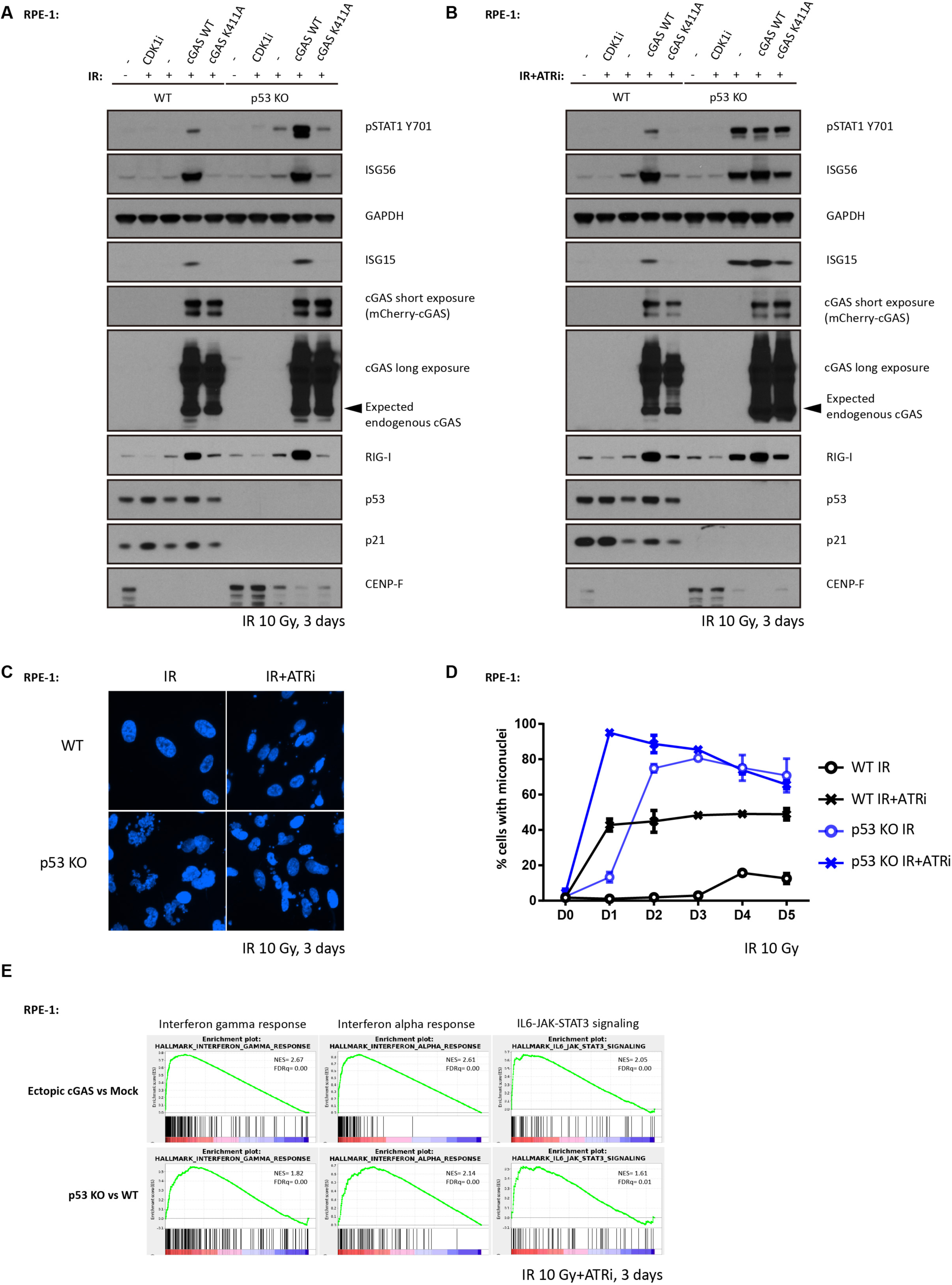
p53 loss cooperates with ATR inhibition activating cGAS-independent inflammatory signaling in RPE-1 cells. (A and B) WT cells or p53 KO cells were irradiated with 10 Gy and cultured in medium with (A) or without ATR inhibitor (B) followed by collection for western blot analysis 3 days later. (C and D) WT cells or p53 KO cells were irradiated with 10 Gy and maintained in medium with or without ATR inhibitor before fixation at indicated time. Cells with micronuclei were qualified (D) and representative images for cells with micronuclei 3 days post IR were showed in (C). (E) RNA-seq data of ectopic cGAS vs mock and p53 KO vs WT were interrogated by Gene Set Enrichment Analysis (GSEA) to identify enriched biological pathways in RPE-1 cells 3 days post 10 Gy irradiation in the presence of ATR inhibitor, respectively. Significant GSEA Enrichment Score curves were noted for interferon alpha response, interferon gamma response, and IL6-JAK-STAT3 signaling. In GSEA thumbnails, the green curve represents the enrichment score curve. Genes on the far left (red) correlated with former cells, and genes on the far right (blue) correlated with latter cells. The vertical black lines indicate the position of each gene in the studied gene set. The normalized enrichment score (NES) and false discover rate (FDR) are shown for each pathway. Error bars represent S.E.M.

### RIG-I mediates inflammatory signaling in RPE-1 p53 KO cells with ATRi treatment

cGAS independence following combined disruption of G1/S and G2/M checkpoints raises the possibility that non-DNA sensing pattern-recognition receptor(s) underlie inflammatory signaling in this setting. RIG-I has been reported to mediate inflammatory signaling to several different stimuli and has emerged as a candidate for cGAS-independent responses (Ablasser et al., 2009; Chiu et al., 2009; Nabet et al., 2017). RIG-I KO single clones were generated using RPE-1 p53 KO cell line and its functional deficiency was confirmed using the RIG-I specific agonist 5’ppp-dsRNA (Figure S5A). Importantly, STING activity was unaffected in RPE-1 p53 and RIG-I double KO clones. 2’3’-cGAMP activated STING-mediated increases in pSTAT1, ISG56 and ISG15 (Figure S5B). RIG-I loss in more than 10 independent RPE-1 p53KO clones compromised the inflammatory signals pSTAT1, ISG56 and ISG15 to varying degrees in response to combined IR and ATRi treatment (Figure 6A and S5C). Consistently, RIG-I was also required for ISG expression in the human colon cancer cell line HCT116 p53-/-, (Figure S5D), which like RPE-1 cells, lacks detectable cGAS expression. Importantly, downstream effectors of the RIG-I pathway were activated as evidenced by the presence of MAVS aggregates (Figure 6B) in RPE-1 p53 KO cells after combined treatment with IR and ATRi. This confirms RNA sensing pathway activation in response to DNA damage as opposed to an indirect effect of RIG-I on another pattern recognition receptor mechanism. RIG-I has been reported to sense RNA transcripts synthesized by RNA polymerase III (Pol III) on cytoplasmic poly (dA:dT) DNA templates (Ablasser et al., 2009; Chiu et al., 2009). Therefore, we tested whether RIG-I mediated inflammatory signaling in RPE-1 p53 KO cells also involve Pol III. We observe Pol III localized to unruptured micronuclei, but not ruptured micronuclei as measured by eGFP-BAF signal (Denais et al., 2016) suggesting that a fraction of micronuclei are Pol III positive (Figure S5E). To explore whether Pol III has a functional role in IR+ATRi induced inflammatory signaling we tested if addition of Pol III inhibitor (Pol IIIi) could block these signals. Consistent with previous reports, Pol IIIi significantly impaired inflammatory signaling stimulated by poly (dA:dT) DNA (Figure S5F). In contrast, Pol IIIi did not affect the inflammatory signals pSTAT1, ISG56 and ISG15 in IR and ATRi treated cells. Together, these results imply that mediators other than Pol III are responsible for RNA sensing pathway activation in checkpoint disrupted cells.

**Fig 6.**
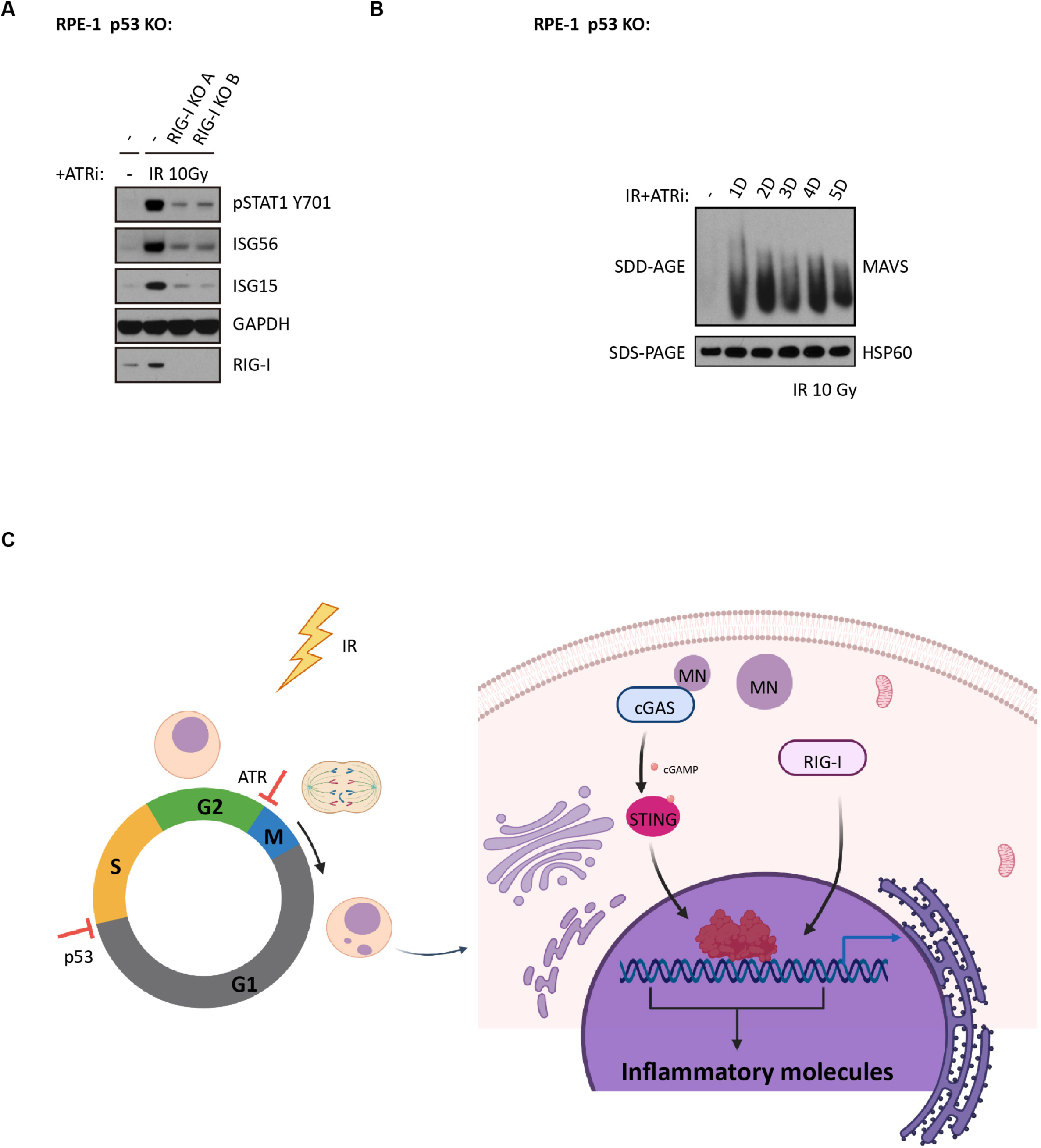
RIG-I mediates inflammatory signaling in RPE-1 p53 KO cells following IR and ATRi treatment. (A) RPE-1 p53 KO cells and RPE-1 p53 RIG-I DKO cells were irradiated with 10 Gy and subjected to culture in medium with ATRi for 3 days before collection for western blot analysis. (B) RPE-1 p53 KO cells were irradiated with 10 Gy and subjected to culture in medium with ATRi for indicated time before crude mitochondria isolation. Equal amounts of isolated crude mitochondria were loaded into 1.5% vertical agarose gels with 0.1% SDS for SDD-AGE analysis. (C) Scheme showing working model of this study. Created with BioRender.

## Discussion

cGAS detects cytosolic dsDNA and triggers production of type I interferons and other inflammatory cytokines and chemokines (Sun et al., 2013). Following IR-induced genotoxic stress, cells forming micronuclei stimulate cGAS-STING mediated innate immune activation and anti-tumor immunity. Cell cycle checkpoints appear to play critical roles in this process since abolishment of mitotic progression compromises micronuclei formation and inflammatory signaling (Harding et al., 2017; Mackenzie et al., 2017). Tumors, however, develop distinct mechanisms of immune evasion, including suppressed expression of pattern recognition receptors cGAS and STING to escape the immune surveillance (Beatty and Gladney, 2015; Ishizuka et al., 2019; Steven and Seliger, 2018; Xia et al., 2016).

This study demonstrates that G2/M checkpoint adaptation links the activation of various inflammatory signaling pathways to anti-tumor immune responses following DNA damage. While abrogation of the G2/M checkpoint facilitates micronuclei formation and inflammatory signaling in cells with intact DNA repair, it was not sufficient in cells that resided at the G2/M boundary for prolonged periods due to excessive, unrepaired DSBs. This scenario required combined disruption of both the G1/S and G2/M checkpoints. Unexpectedly, loss of p53 together with ATR inhibition upregulated inflammatory signaling in RPE-1 cells with undetectable cGAS, suggesting that cell cycle checkpoints coordinate to limit signaling through multiple pattern recognition receptor pathways and conversion of intrinsic tumor stress signals into anti-tumor immune responses. Our findings reveal the cytosolic RNA sensor RIG-I as an additional component of this response (Figure 6C). This is also consistent with our original report that cGAS or STING loss did not eliminate IR induced inflammatory stimulated gene expression (Harding et al. 2017). In further support, combined IR and ATRi significantly increased interferon production and immune infiltration *in vivo* compared to IR alone in TC-1 tumors with p53-inactivation due to human papilloma virus (Dillon et al., 2019). Together with our findings, this implies that combined loss of G1/S and G2/M checkpoints can enhance immune infiltration and anti-tumor responses.

How checkpoint proteins p53 and ATR cooperate to restrict inflammatory signaling warrants further consideration. A potential explanation is that their combined loss increases micronuclei formation and other structures that may attract cGAS. Our data further supports that entry into S-phase and progression through a second round of mitosis enhances micronuclei formation (Figure S2). Disruption of both G1/S and G2/M checkpoints by combined p53 loss and ATR inhibition would augment this accelerated cell proliferation and cytoplasmic DNA accumulation. Interestingly, micronuclei passage through mitosis and chromosome shattering followed by reassembly in the subsequent is thought to occur during the process of chromothripsis (Umbreit et al., 2020; Zhang et al., 2015). Such catastrophic chromosome breakage is predicted to not only promote large-scale genomic rearrangements, but also increase the number of acentric fragments that could missegregate into the cytoplasm and augment inflammatory signaling.

The role of checkpoint proteins p53 and ATR in restricting RIG-I mediated inflammatory signaling is less obvious. Notably, this response also required progression through mitosis but was not RNA Pol III dependent. An alternative possibility is that these cells experience chromatin alterations that drive changes in transcription globally. Reactivation of endogenous retroviral elements following methyltransferase inhibitors are capable of activating RNA sensing pathways and similar activation of endogenous RNA elements have been observed after IR (Chiappinelli et al., 2015; Ranoa et al., 2016; Roulois et al., 2015; Rudin and Thompson, 2001). Interestingly, p53 loss increases derepression of Short Interspersed Nuclear Elements (SINE) after treatment of DSB inducing agents (Hagan and Rudin, 2007). Whether this interplay occurs following IR treatment is an important avenue for future studies and may suggest additional routes to activate anti-tumor immune responses in p53 mutant cancers.

An additional finding in this study is the complex relationship between intrinsic DNA repair capacity and activation of anti-tumor immunity. Loss of c-NHEJ prevented anti-tumor immune responses in syngeneic melanoma models treated with radiotherapy and immune checkpoint blockade. Although c-NHEJ components DNA-PKcs and KU have been reported as DNA sensors for innate immunity (Burleigh et al., 2020; Ferguson et al., 2012; Zhang et al., 2011), we observe similar reductions in cells that have lost end ligation factors XRCC4 and Lig4, (Graham et al., 2016). XRCC4 and Lig4 are not required for DSB end recognition or signaling, making them unlikely candidates for pattern recognition. We propose the effects of c-NHEJ loss are secondary to excessive unrepaired DSBs and persistent cell cycle arrest. Consistent with this assertion, low IR-doses induce slightly higher micronuclei formation and inflammatory signaling in XRCC4 KO cells as compared with high IR-doses. They also predict that lower doses of DNA damaging agents may be necessary to maximally recruit anti-tumor immune responses in the setting of DNA repair deficient cancers. Additional reports also show that fractionated “low” doses do not induce expression of the cytoplasmic nuclease TREX1 to the extent observed for single high doses of IR (Vanpouille-Box et al., 2017). It remains unclear however if this contributed to our observations. Taken together, our work reveals the suppressive role of cell cycle checkpoints in DNA damage induced inflammatory signaling. These data suggest potential approaches to stratify patients based on tumor genotypes and alternative strategies to activate inflammatory signaling in tumors that harbor resistance mechanisms.

## Acknowledgements

We thank all members of the Greenberg laboratory for critical discussion. We are grateful to D Durocher (Lunenfeld-Tanenbaum Research Institute) for providing hTERT RPE-1 cells and hTERT RPE-1 p53 KO cells, and to B. Vogelstein (Johns Hopkins University Medical School) for providing HCT116 p53 KO cells. This work was supported by NIH grants GM101149 and CA17494 and a V Foundation Team Convergence Award (to RAG), who is also supported by funds from the Penn Center for Genome Integrity and Basser Center for BRCA. JC was supported by the Michael Brown Penn-GSK Postdoctoral Fellowship.

---

**Fig S1.**
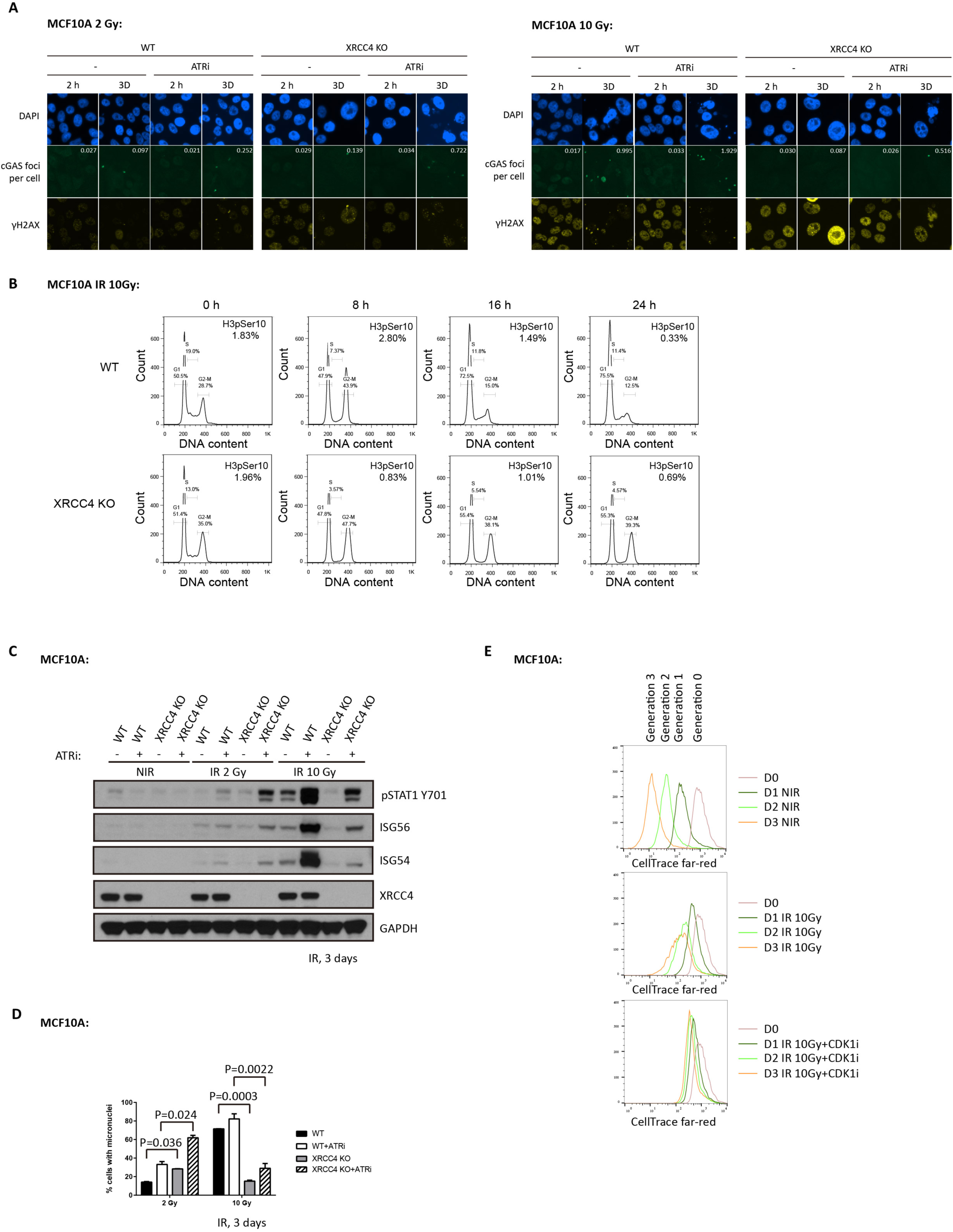
Low dose irradiation allows micronuclei formation and inflammatory signaling in c-NHEJ deficient MCF10A cells. (A) WT cells or XRCC4 KO cells were irradiated with 2 Gy or 10 Gy, and then cultured with or without ATRi for indicated time before fixation for immunofluorescence staining. (B) WT cells or XRCC4 KO cells were irradiated with 10 Gy followed by time course collection for cell cycle and mitotic population analysis using flow cytometry. (C and D) WT cells or XRCC4 KO cells were treated with 2 Gy or 10 Gy IR, and then maintained for 3 days with or without ATRi before western blot analysis (C) or qualification of cells with micronuclei (D). (E) MCF10A cells were resuspended in medium containing 1 uM CellTrace far-red dye (Thermo) for 20 mins and subsequently washed and seeded in dye-free medium. The next day, cells were either left untreated (NIR, top panel) or treated with 10 Gy (middle panel) or 10 Gy+CDK1i (bottom panel), followed by collection for CellTrace far red signal analysis using flowcytometry. Reductions in dye content over time indicate cell division due to dye dilution. Non-irradiated (NIR) cells undergo three divisions whereas irradiated controls perform 1-2 divisions. CDK1i prevents cell division. Error bars represent S.E.M.

**Fig S2.**
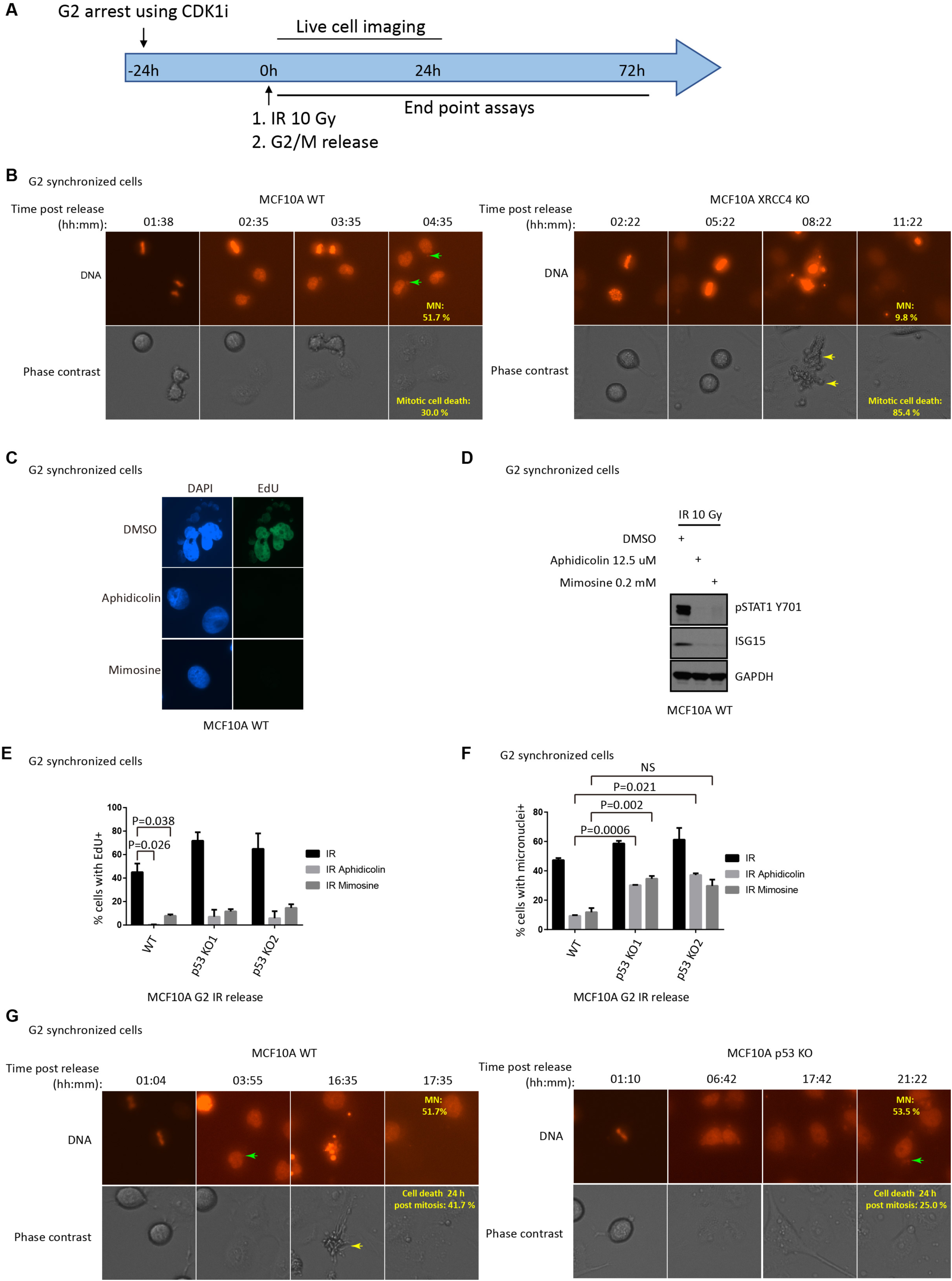
Increased micronuclei formation in MCF10A p53 KO cells after IR. (A) Scheme showing working flow of live cell imaging (B and G) and end point assays (C, D, E and F). (B) G2 arrested MCF10A WT cells and MCF10A XRCC4 KO cells were irradiated with 10 Gy and then released from G2 phase followed by live cell imaging to monitor micronuclei formation. siR-DNA was used to visualize DNA in live cells. Micronuclei were recognized as positive siR-DNA staining outside of primary nuclei. Mitotic cell death was identified as anaphase cells broke into small pieces, detached from dish, and floated in medium under both phase contrast and siR-DNA staining channels. (C and D) G2 arrested MCF10A cells were irradiated and then released but blocked before next S phase to probe micronuclei formation in the presence of EdU (C) and inflammatory signaling (D). (E and F) MCF10A WT cells and MCF10A p53 KO cells were treated as in (C) for quantification of cells with EdU incorporation (E) and cells with micronuclei (F). (G) MCF10A WT cells and MCF10A p53 KO cells were treated as in (B) to monitor cell death after mitosis in the following 24 hours. Cell death post mitosis was qualified when cells broke into small pieces and detached from the slide following mitotic progression with micronuclei under both phase contrast and siR-DNA staining channels. Error bars represent S.E.M.

**Fig S3.**
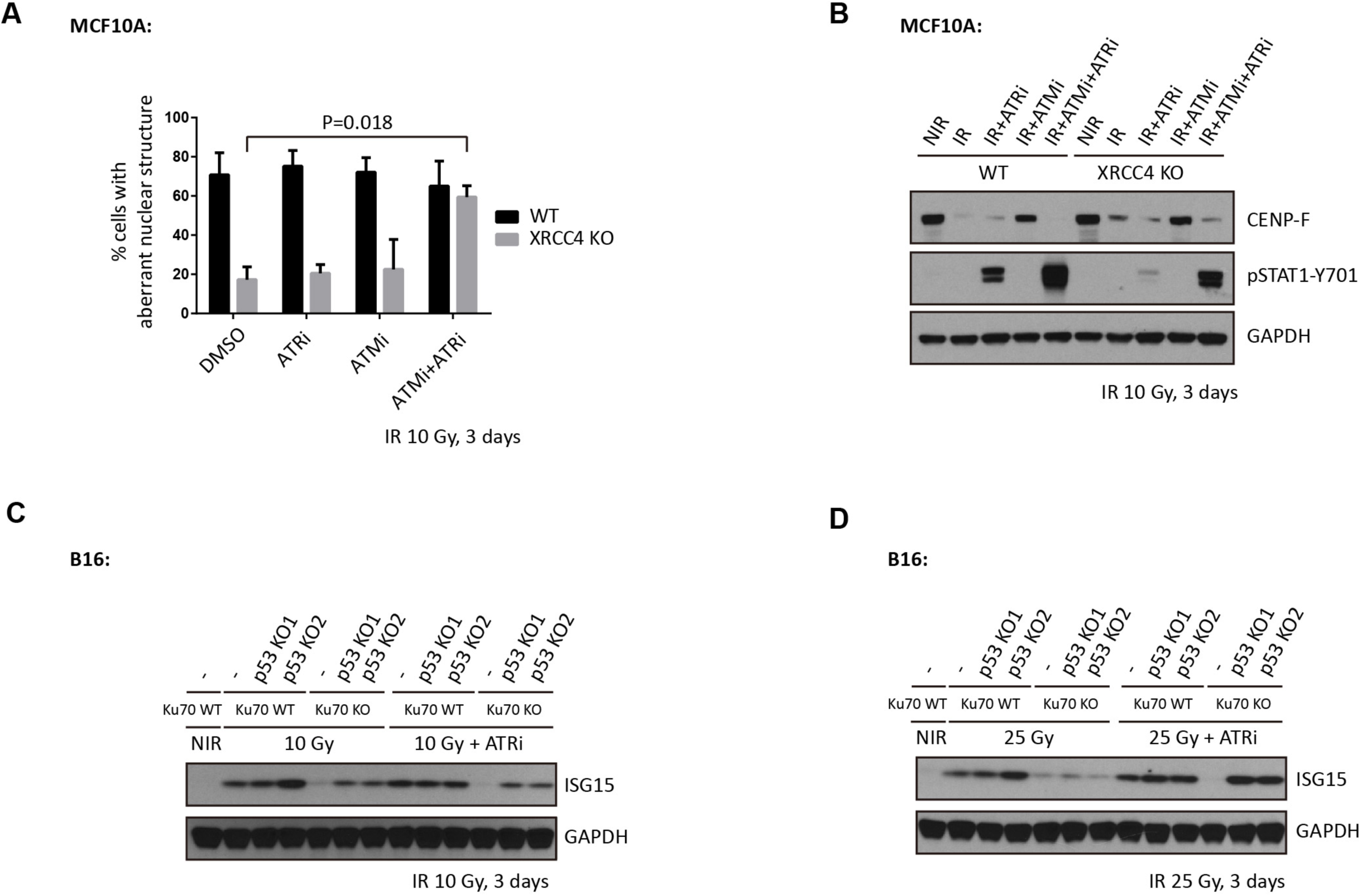
Restored inflammatory signaling after disruption of G1/S and G2/M checkpoints in c-NHEJ deficient cells. (A and B) MCF10A WT cells or XRCC4 KO cells were irradiated with 10 Gy and then maintained in medium with indicated inhibitors for 3 days before qualification for cells with micronuclei (A) or western blot analysis (B). (C and D) B16 WT cells and B16 cells with p53 KO or/and Ku70 KO were subjected to 10 Gy IR (C) or 25 Gy IR (D) treatment. Cells were then maintained in medium with or without ATR inhibitor for 3 days before western blot analysis. Error bars represent S.E.M.

**Fig S4.**
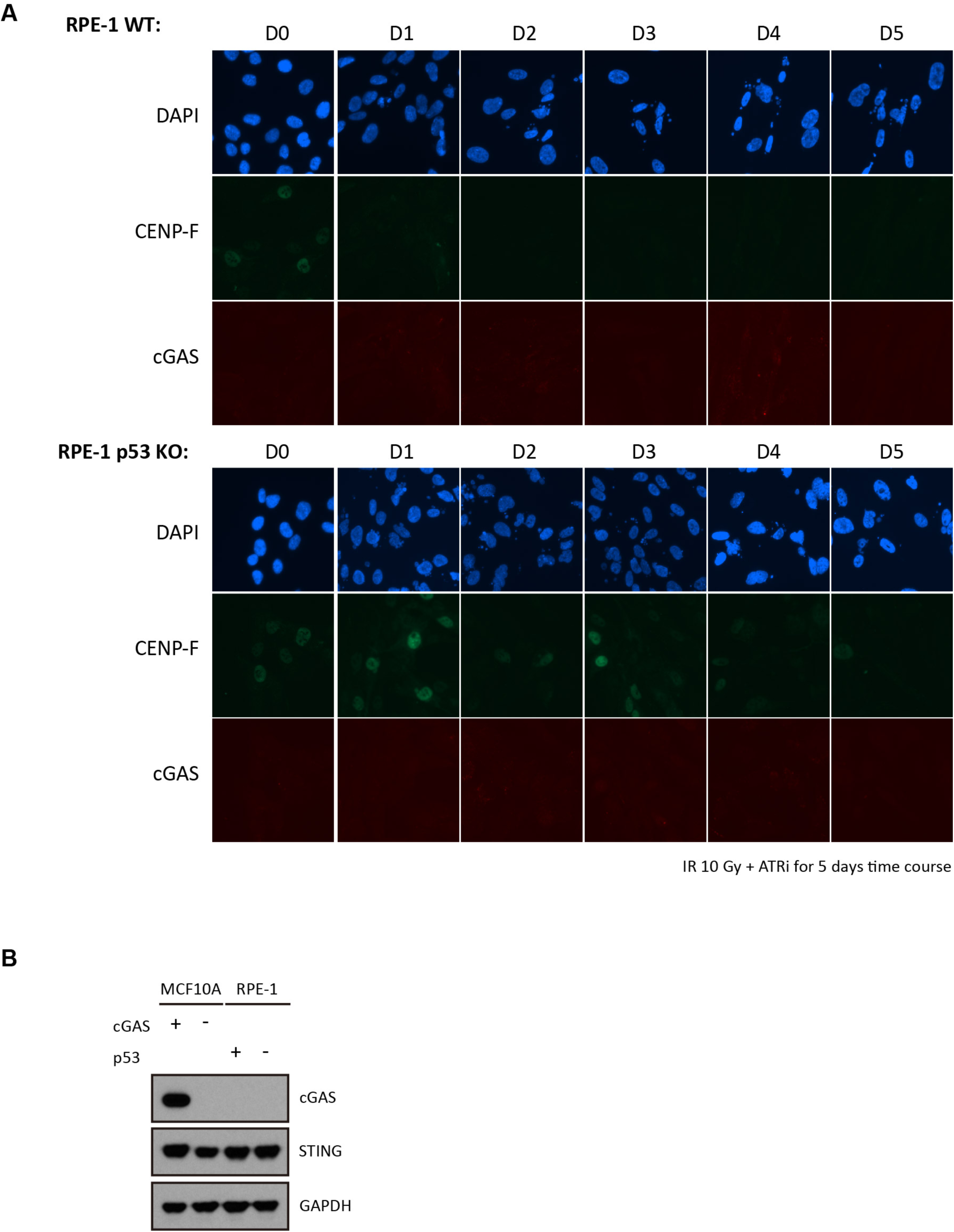
Undetectable cGAS expression in RPE-1 cells. (A) RPE-1 WT cells and RPE-1 p53 KO cells were irradiated with 10 Gy and subjected to culture in medium with ATRi. Cells were then fixed at indicated time for immunofluorescence staining. (B) MCF10A cells and RPE-1 cells were collected to probe cGAS expression using western blot analysis.

**Fig S5.**
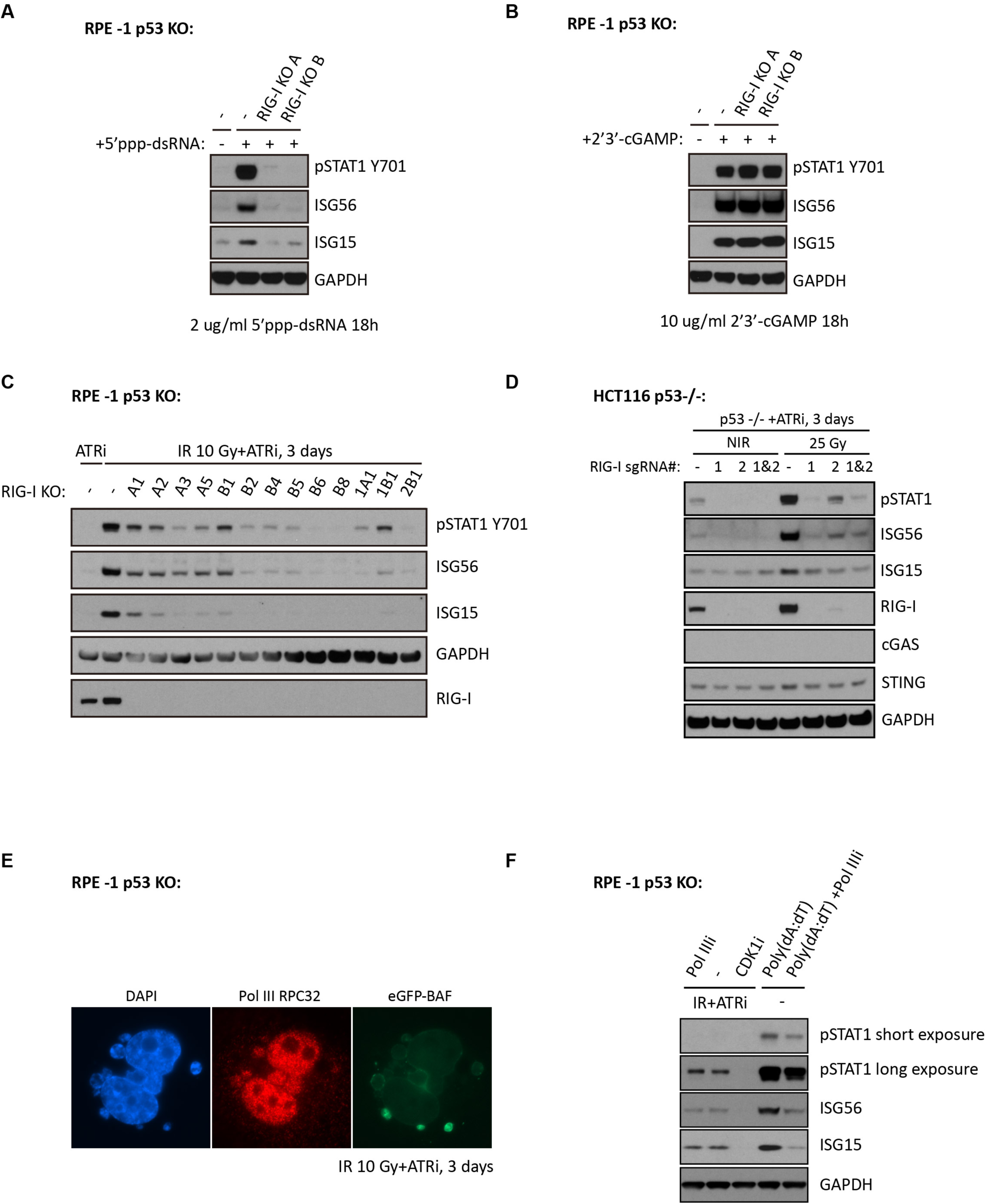
Pol III independent RIG-I activation in RPE-1 p53 KO cells following IR plus ATRi treatments. (A and B) RPE-1 p53 KO cells and RPE-1 p53 RIG-I DKO cells were transfected with RIG-I agonist (A) or STING agonist (B) and then collected 18 hours later for western blot analysis. (C) RPE-1 p53 KO cells and RPE-1 p53 RIG-I DKO single clone cells were irradiated with 10 Gy and subjected to culture in medium with ATRi for 3 days before collection for western blot analysis. (D) RIG-I depleted HCT116 p53-/-pool cells were achieved by sequential lentiviral infections of Cas9 and sgRNAs targeting RIG-I (#1, #2, or #1&2). Parental HCT116 p53-/-cells and HCT116 p53-/-cells with RIG-I depletion were left untreated (NIR) or treated with 25 Gy followed by culture in medium with ATRi for 3 days before collection for western blot analysis. (E) RPE-1 p53 KO cells expressing ectopic eGFP tagged BAF were irradiated with 10 Gy and incubated in medium containing ATRi for 3 days before fixation for immunofluorescence staining. 10 Gy irradiated RPE-1 p53 KO cells were maintained in medium with 2.5 μM ATRi, 2.5 μM ATRi+ 40 μM Pol IIIi or 2.5 μM ATRi+ 9 μM CDK1i for 3 days before collection for western blot analysis. Cells transfected with poly(dA:dT) at a final concentration of 0.5 μg/ml were cultured in the absence or presence of 40 μM Pol IIIi for 1 day before western blot analysis.

## Methods

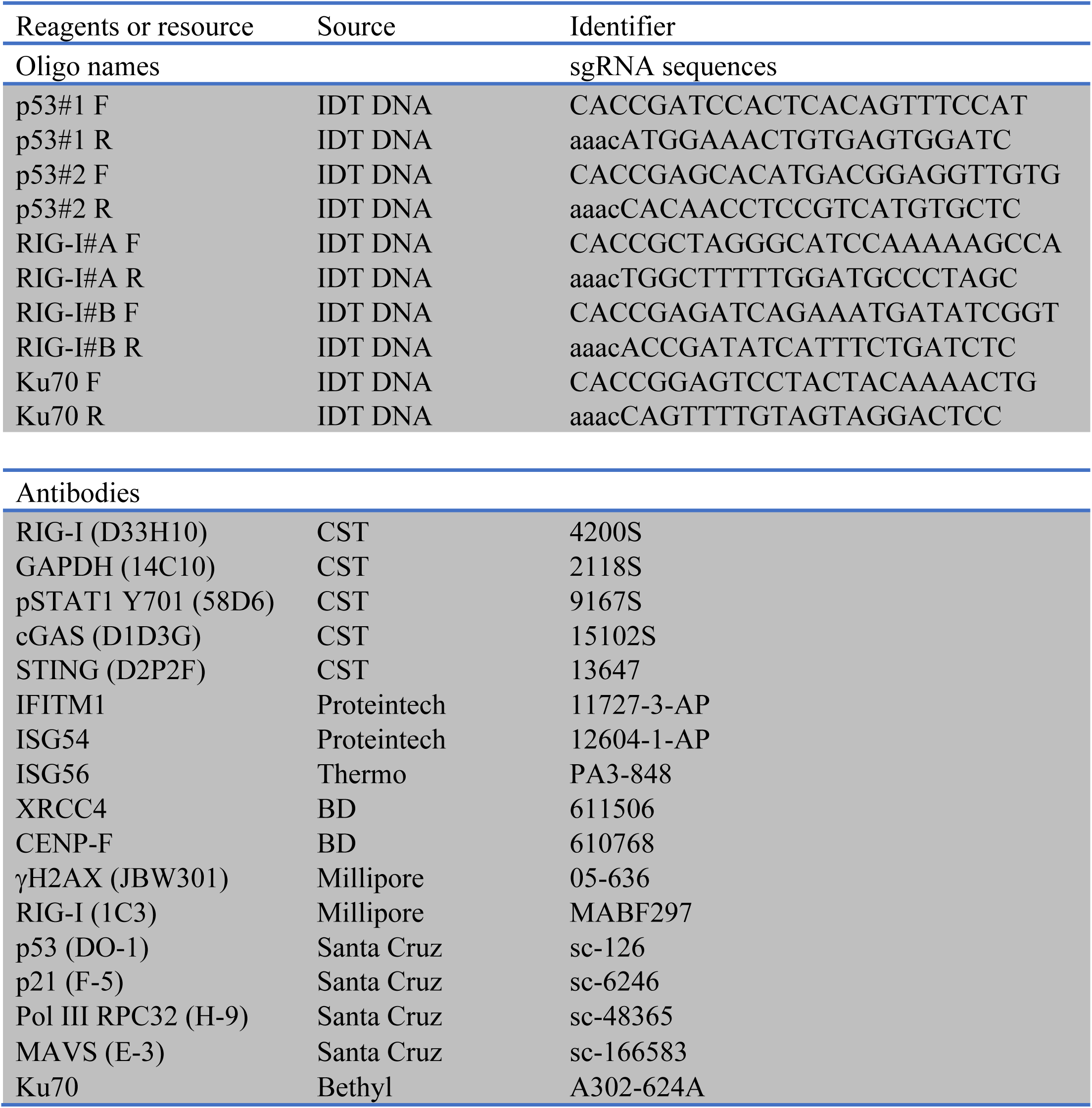

### Cell lines and tissue culture

MCF10A cells were obtained from ATCC and cultured in DMEM/F-12 media with 5% horse serum (Thermo Fisher Scientific), 20 ng/ml human EGF (Peprotech), 0.5 mg/ml hydrocortisone, 100 ng/ml cholera toxin and 10 μg/ml recombinant human insulin (Sigma). MCF10A I-PpoI cells and MCF10A AsiSI cells were previously described (Harding et al., 2017). hTERT RPE-1 cells and hTERT RPE-1 p53 KO cells were kindly provided by Dr. Dan Durocher (Lunenfeld-Tanenbaum Research Institute) and cultured in DMEM media with 10% bovine calf serum (Hyclone). B16-F10 cells were purchased from ATCC and cultured in DMEM with 10% bovine calf serum (Hyclone). HCT116 p53-/-cells (obtained from B Vogelstein) were cultured in McCoy’s 5a media with 10% bovine calf serum (Hyclone). All cells were maintained with penicillin and streptomycin (Thermo Fisher Scientific).

### Plasmid construction

pLVX-mCherry-cGAS was generated as previously described (Harding et al., 2017). pLVX-mCherry-cGAS K411A was generated by site-directed mutagenesis. LentiGuide-puro targeting p53 and RIG-I was generated by inserting targeting duplex oligos into LentiGuide-puro at BsmBI site.

### Irradiation and drug treatments

Cells were seeded at densities of 40%-50% at the time of treatment. Cells were irradiated using a Cs-137 Gammacell irradiator (Nordion) with ∼0.76 Gy/min. Inhibitors were administrated 1 h prior to irradiation and maintained till collection unless otherwise stated. Media with or without inhibitors were changed every 2 days after treatment. Medium containing Pol IIIi were changed every day. Inhibitors were used at following concentrations: CDK1i (9 μM, RO-3306, Selleck Chemical), ATRi (2.5 μM, VE-821, Selleck Chemical), ATMi (10μM, Ku55933, Selleck Chemicals), and Pol IIIi (40 μM, ML-60218, Millipore). Agonists were transfected using Avalanche-Omni (EZT-OMNI-1, EZ Biosystems) and incubated for 18 hours before collection. Agonist were used as following concentrations: 5’ppp-dsRNA (2 μg/ml, tlrl-3prna, Invivogen) and 2’3’-cGAMP (10 μg/ml, tlrl-nacga23, Invivogen). For AsiSI or I-PpoI nuclease induced DSBs, cells were treated with Shield-1 (1 μM, AOB1848, Aobious) and 4-OHT (2 μM, H7904, Sigma) for 5 hours before washing with PBS and adding back media.

### CRISPR–Cas9 knockout

Lentiviruses of LentiCas9-blast and LentiGuide-Puro (gifts from F. Zhang, Addgene #52962, Addgene#52963) were produced and concentrated as previously described (Kutner et al., 2009). Cells were infected with lentiviruses overnight in the presence of 8 ug/ml polybrene. Cells infected with LentiCas9 were subjected to 10μg/ml blasticidin (Invivogen) until control cells without infection were all killed. Knockout clones were generated by infecting cells with lentivirus using lentiGuide-Puro followed by selection in 2 μg/ml puromycin (Sigma) for 2 days. Single colonies were achieved by seeding one cell per 96-well. p53 knockout clones were validated by western blotting using anti-p53 antibody (DO-1, santa cruz). RIG-I knockout clones were validated by western blotting using anti-RIG-I antibody (D33H10, CST).

B16-F10 Ku70 knockout clones were created as previously described (Benci et al., 2016). Ku70 knockout clones were validated by western blotting using anti-Ku70 antibody (A302-624A, bethyl antibody)

### Western blotting

Western blotting was performed using standard methods. In brief, cells were collected by trypsinization, washed in PBS and lysed in NETN buffer (150 mM NaCl, 1% NP-40 alternative, 50 mM Tris pH 7.4) with turbo nuclease (Accelagen) in the presence of 1 mM MgCl2, protease inhibitor cocktail (Roche) and phosphatase inhibitor cocktail (5 mM sodium fluoride, 1 mM sodium orthovanadate, 1 mM sodium pyrophosphate decahydrate, 1 mM β-glycerophophate) on ice for 30 min. Protein was quantified using Bradford method. Equal amount of protein was separated on 4-12% Bis-Tris gels (Thermo Fisher Scientific) using MOPS/MES buffer and transferred to 0.2 µM Amersham Protran Premium Nitrocellulose Western Blotting Membranes (10600004, GE) at 350 mA for 2 h in ice cold transfer buffer (20% methanol, 191 mM glycine, 25 mM tris-base, 0.1% SDS, pH 8.3). Membrane were blocked in 3% non-fat milk in PBST (0.1% Tween 20) and incubated overnight at 4°C in primary antibody diluted with PBS containing 1% BSA. Membrane were washed in PBST for 20 min and incubated at room temperature for 2 h in secondary antibody (Amersham ECL HRP, GE) diluted with PBST containing 3% non-fat milk. Blots were developed using immobilon forte western HRP substrate (WBLUF0100, Millipore).

### *In vivo* MAVS aggregation assay

*In vivo* MAVS aggregation assay was performed according to a published protocol (Hou et al., 2011). Briefly, crude mitochondria isolated using mitochondria isolation kit (89874, Thermo) were resuspended in 1x sample buffer (0.5 x TBE, 10% glycerol, 2% SDS) supplemented with protease inhibitor cocktail (Roche) and phosphatase inhibitor cocktail (5 mM sodium fluoride, 1 mM sodium orthovanadate, 1 mM sodium pyrophosphate decahydrate, 1 mM β-glycerophophate) followed by protein concentration quantification using BCA method. Equal amounts of crude mitochondria were mixed with 1/10 volume gel loading dye (7025, NEB) and subjected to Semi-Denaturing Detergent Agarose Gel Electrophoresis (SDD-AGE). Samples were loaded onto a homemade vertical 1.5% agarose gel prepared using Novex empty gel cassettes (NC2015, Thermo), empty gel cassette combs (NC3515, Thermo) and Bio-Rad agarose (1613101, BioRad). After electrophoresis in the running buffer (1 x TBE and 0.1% SDS) for 40 min with a constant voltage of 100 V at 4°C, the proteins were transferred to 0.2 µM Amersham Protran Premium Nitrocellulose Western Blotting Membranes (10600004, GE) at 400 mA for 3 h in ice cold transfer buffer (10% methanol, 191 mM glycine, 25 mM tris-base, 0.1% SDS, pH 8.3) followed by immunoblotting.

### Immunofluorescence

Cells were seeded onto coverslips in 24-well plate one day before treatment and fixed with 3% paraformaldehyde (PFA). Cells were then washed with PBS, permeabilized in 0.5% Triton X-100 solution for 1 min at room temperature and processed for immunostaining using the indicated antibodies. Coverslips were mounted in VECTASHIELD Antifade Mounting Medium with DAPI (H-1200-10, Vector Labs). Images were taken using a Nikon Eclips 80i microscope with a Coolsnap Myo camera (Photometrics) and Nikon NIS-Elements software. Images were prepared using FIJI (NIH). Cells with aberrant nuclear structure in cytoplasm were visualized by DAPI staining and counted manually.

### Live-cell imaging

MCF10A cells were seeded onto falcon 24-well plates (08-772-1H, Fisher) and cultured in medium as described above in which DMEM/F-12 base medium was replaced by 1:1 mixed FluoroBrite DMEM (A1896701, Thermo) and phenol red free F12 medium (HFL05-500ML, Caisson Labs). CDK1i arrested cells were irradiated on ice, washed, replaced with medium containing 0.5 μM siR-DNA (CY-SC007, Cytoskeleton), a cell permeable far-red probe for DNA, and then immediately subjected to live-cell imaging using EVOS FL Auto Imaging System (Thermo). Imaging was carried out in a 37 °C humidified chamber equilibrated with 5% CO2 using a 20x air objective. Images were acquired every 20 min for 20-24 h.

### Flow cytometry

Cells were dissociated into single cells with trypsin, washed once in PBS, resuspended in 300 μl PBS, and fixed by drop-wise addition of 700 μl pre-chilled (−20°C) 100% ethanol. After fixation at 4 °C or storage at -20 °C, cells were washed once with 6 ml 1% BSA/PBS and pelleted by centrifuge. Cell pellets were loosed and resuspended by flick in 100 μl 1% BSA/PBS with 1 μl Alexa Fluor 488 conjugated phospho-Histone H3 (Ser10) antibody (DC28, Cell Signaling). After incubation at room temperature for 1 hour with occasional flick, cells were washed once with 6 ml 1% BSA/PBS and resuspended in PBS containing 10 μg/ml propidium iodide (Santa Cruz) and 100 μg/ml RNase A (Qiagen). Flow cytometry was performed on a BD FACSCalibur and analyzed with FlowJo software. Single cell population and G1/S/G2 population were manually gated.

### Modified RadVax procedure

Experiment was performed as previously described (Harding et al., 2017). In brief, five-to seven-week-old female C57BL/6 mice were obtained from Charles River and maintained under pathogen-free conditions. Mice were divided randomly into cages upon arrival and were randomly injected and measured without blinding. All animal experiments were performed according to protocols approved by the Institute of Animal Care and Use Committee of the University of Pennsylvania (IACUC). The minimal number of animals was used based on prior experience to yield consistent measurements. On the day before the experiment, B16-F10 cells were treated *in vitro* with 10 Gy of ionizing radiation and cultured with or without CDK1i. CDK1i was added 1h before irradiation and maintained until cell isolation and injection on day 2. On day 0 untreated B16-F10 cells (5× 10^4^) in 50μl of PBS were mixed with an equal volume of Matrigel (356237, Corning) and injected into the right flank. On day 2, 5×10^5^ IR treated cells including B16-F10 parental cells, parental cells with CDK1i treatment and Ku70 knockout cells were mixed with Matrigel and injected on the opposite flank. On days 5, 8 and 11 anti-CTLA4 antibody (9H10; BioXCell) was administered interperitoneally at 200 μg per mouse. Volumes were measured using calipers starting at day 11 and calculated using the formula *l*× *w*2× 0.52, where *l* is the longest dimension and *w* is perpendicular to *l*. Animals were euthanatized when either tumor reached 1.5 cm in the largest dimension according to IACUC guidelines.

### RNA purification, RNA-seq library preparation and analysis

Total RNA for RNA-seq library was purified using miRNeasy Mini Kit (217004, Qiagen) and validated on a agilent RNA 6000 nano chip (Agilent Technologies) with a RNA integrity number (RIN) > 8. RNA-seq libraries were prepared using the TruSeq Stranded mRNA Library Prep (20020594, Illumina), TruSeq RNA Single Indexes Set A (20020492, Illumina) and TruSeq RNA Single Indexes Set B (20020493, Illumina) according to standard Illumina library preparation procedure. In brief, 0.5 μg of purified RNA was poly-A selected and fragmented followed by first and second strand cDNA synthesis. Double-stranded cDNA was processed from end-repair to PCR amplification according to library construction steps. Libraries were purified using AMPure XP beads (A63880, Beckman Coulter) and validated for appropriate size on a 2100 Bioanalyzer High Sensitivity DNA chip (Agilent Technologies, Inc.). The DNA library was quantitated using Qubit (Thermo Fisher) and normalized to a concentration of 4 nM prior to pooling. Libraries were pooled in an equimolar fashion and diluted to a final concentration of 2 pM. Library pools were clustered and run on a Nextseq500 platform with single-end reads of 75 bases (20024906, Illumina), according to the manufacturer’s recommended protocol (Illumina Inc.). Differential gene expression analysis between the target and reference sets of treatments were determined using DESeq2 (Love et al., 2014) (https://bioconductor.org/packages/release/bioc/html/DESeq.html). Gene Set Enrichment Analysis (GSEA) was performed with the Molecular Signatures Database (MSigDB) hallmark gene set (Liberzon et al., 2015; Subramanian et al., 2005). Both DESeq2 and MSigDB are provided by the Broad Institute.

## Statistical Analysis

All statistical analyses were carried out using GraphPad Prism. For quantitative data, data was presented as mean ± S.E.M. and statistical significance was analyzed using two-tailed unpaired Student’s t test.

